# Nyctinastic leaf folding mimic reduces herbivory by *Chromacris trogon* grasshoppers (Orthoptera:Romaleidae)

**DOI:** 10.1101/2023.06.16.545391

**Authors:** Aidan B. Bell

## Abstract

*Arachis pintoi* (Fabaceae) is a common relative of the cultivated peanut, and folds its four leaflets up to look like one at night. The adaptive significance of this behavior (foliar nyctinasty) is unknown. To test the hypothesis that leaflet folding alone can deter herbivores, a leaf preference experiment was performed on *Chromacris trogon* grasshoppers. Small oval cutouts were made from leaves of the grasshopper’s preferred food source, *Iochroma arborescens* (Solanaceae), and were combined with small pieces of tape and dry grass to construct artificial leaves resembling the day and night form of *A. pintoi*. In the experiment, groups of three grasshoppers were starved for 24 hours and then placed in petri dishes containing one closed and one open artificial leaf. After 30 six-hour trials, the average herbivory of open leaves was 12.3%, while closed leaves was 5.2% (p = 0.00145), indicating a significant preference for open leaves. This suggests that the folded configuration of *A. pintoi* leaves can be a defense against herbivory.

## INTRODUCTION

*Arachis pintoi* (pinto peanut) is native to Brazil, but is now found in many tropical and subtropical regions, including Australia, South, Central, and North America, and South-East Asia. It is commonly used as a pasture plant or cover crop (Das et al. 2020). Like many other legumes, it exhibits foliar nyctinasty, folding its four leaflets up to look like one at night (see Fig. 1). The adaptive significance of the foliar nyctinasty of *A. pintoi* is unknown. In a review of foliar nyctinasty (abbreviated as FN), Minorsky (2019) defined FN as a plant behavior characterized by a pronounced daily variation in leaf or leaflet orientation. FN is a surprisingly widespread phenomenon, existing in over 200 plant genera across 38 families (Minorsky, 2019). FN is found in plants adapted to xeric, mesic or aquatic environments, tropical or temperate zones, and exists in forbs, shrubs and tall trees (Minorsky, 2019). There is considerable diversity in the biochemical mechanisms underlying motion in FN (Ueda, Shigemori, & Sata, 2000), which can involve pulvinar and non-pulvinar action (Wetherell, 1990) Due to the wide taxonomic distribution of FN and the variety in biochemical mechanisms that cause it, it is thought that FN has convergently evolved many times (Minorsky, 2019). Moreover, as guard cell swelling incurs a bioenergetic cost (Assmann & Zeiger, 1987), it is reasonable to posit that nyctinastic movements also entail an energetic cost. The energetic cost of FN, combined with its probable convergent evolution, suggest that there is likely some adaptive significance of FN (Minorsky, 2019).

**Figure 1.**
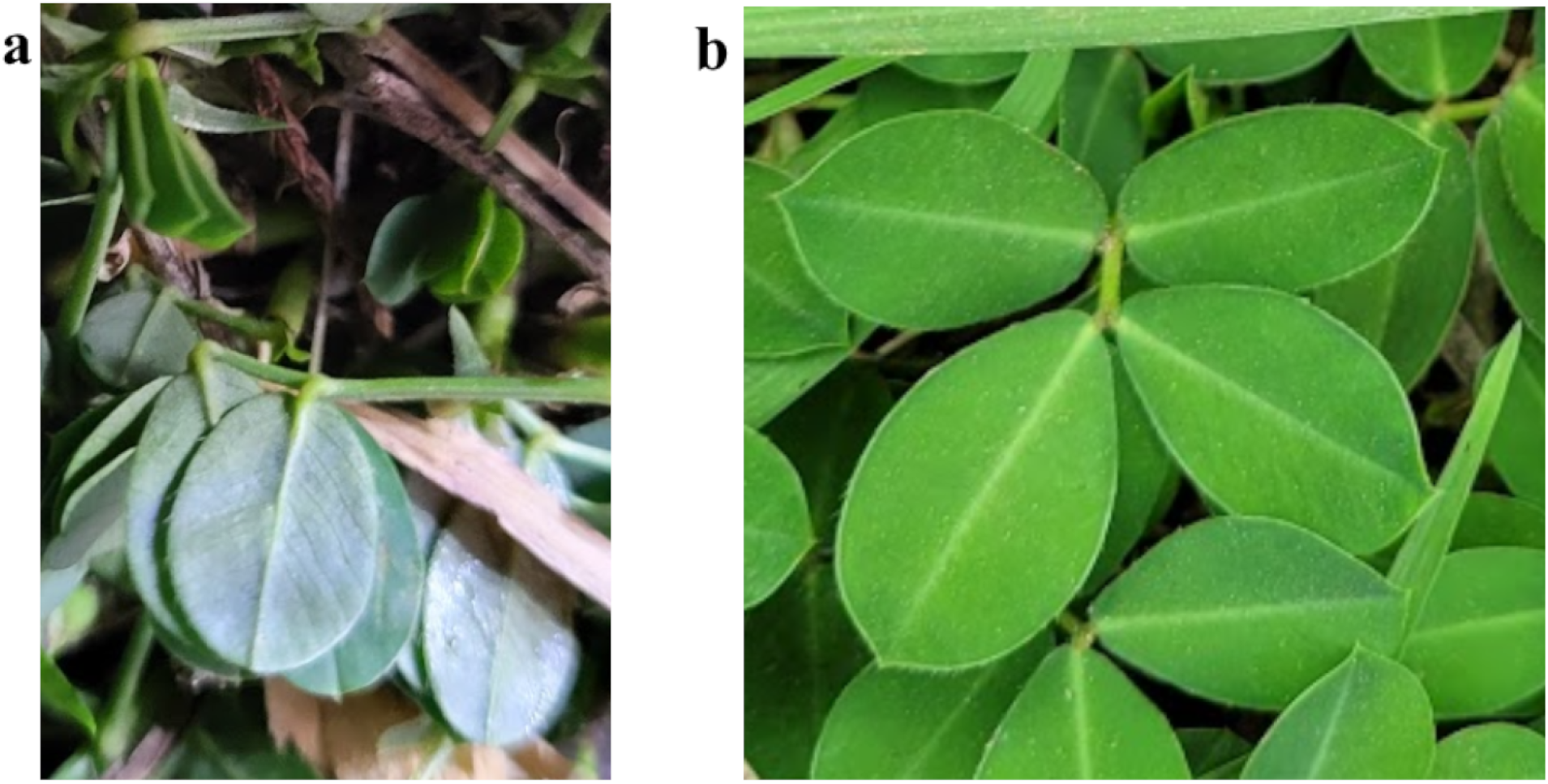
**a** *A. pintoi* leaves folded during the night and **b** open during the day. *A. pintoi* also has secondary pulvinus movement at night, moving the entire folded leaf and petiole closer to the ground. The small, folded leaves in the upper left of **a** are new leaves, and stay folded during the day.

According to Minorsky (2019), the prominent hypotheses for the broad adaptive significance of FN are (1) improving temperature relations, (2) assisting in removing surface water, and (3) acting as a defense against herbivory. FN can help plants survive subzero temperatures, and closing of rosette leaves has been shown to prevent freezing in *Espeletia schultzii* (Smith, 1974). However, this explanation fails to account for the widespread occurrence of FN in the Fabaceae family, which evolved in tropical regions devoid of freezing conditions (Herendeen et al., 1992). In *Phaseolus vulgaris*, the common bean (Fabaceae), FN completely stopped within 1 day at 5°C (Hoshizaki & Hamner, 1969), suggesting that at least in some legumes, FN is not an adaptation to deal with extreme cold. Since *A. pintoi* does not experience freezing conditions in its environment, freezing prevention by FN provides no adaptive advantage to *A. pintoi*.

FN has been shown to aid in removing surface liquid (Dean & Smith, 1978; Gitari, 1986), and this is hypothesized to affect plant fitness in a number of ways. Removing surface liquid has been hypothesized to improve fitness by (1) increasing transpiration and thus nutrient uptake (but little to no transpiration happens at night), (2) reducing mineral leaching into surface water, (3) reducing the structural stress of the weight of water, and (4) reducing growth of fungal pathogens and epiphytes that prefer surface liquid on leaves (Smith, 1978). Surface liquid removal hypotheses could explain the FN of *A. pintoi*, but it is important to note that there are many more efficient forms of removing surface moisture such as smooth cuticles, non-horizontal leaf angles, certain forms of pubescence, and drip-tips, that are functional all of the time (Minorsky, 2019). *A. pintoi* is found in inland tropical and subtropical regions, where most rain happens during the late afternoon, not at night when FN happens (Yang & Smith, 2006). Although it could be a purpose of FN in *A. pintoi*, this study will not focus on removal of surface liquid as an explanation of FN in *A. pintoi*. For a more in depth review of these hypotheses, see Minorsky (2019) and Smith (1978).

FN has also been hypothesized to be a defense against herbivory (Minorsky 2019), a hypothesis I attempt to directly test in this study. To my knowledge, there is currently no experimental evidence showing that leaf folding is a defense against herbivory on its own. Leaf folding has been shown to be correlated with herbivore defense in a number of studies, however. The closure of *Mimosa pudica* leaves in response to touch, has been experimentally confirmed to reduce herbivory by grasshoppers and caterpillars in an experiment led by Hagihara and coworkers (2022). Although the folded configuration of *M. pudica* leaves is likely a factor in the reduction of herbivory observed by Hagihara and coworkers (2022), folding cannot be separated from the effect of leaflets rapidly moving in response to stimulation in this study. Jensen and coworkers (2011) found that *Mimosa pudica* leaves stay in the closed configuration longer after stimulation when not stressed by lack of sunlight, even though closing leaves reduces photosynthesis (Hoddinott, 1977). This suggests that the folded leaf configuration confers a benefit greater than the reduced photosynthesis efficiency when there is lots of light. *M. pudica* does expose thorns when leaflets are folded (Eisner, 1981), so thorn exposure could explain *M. pudica* leaves staying folded for extended periods of time.

When discussing leaf folding as a potential herbivore defense, it is important to bring up how many young leaves of plants emerge folded. 22 of the 39 monocot families, and 7 dicot families found in tropical lowland rainforest rain forest have species with leaves folded until they are >50% of the final length (Grubb & Jackson, 2007). Grubb and Jackson (2007) hypothesize that this folding is a defense against herbivory. In a study of five European tall shrub species, Jackson and coworkers (1999) found a lower herbivory rate on new, folded/rolled leaves compared to freshly unfolded leaves in the field. Jackson and coworkers (1999) show leaf folding to be correlated with low attractiveness to herbivores, proposing that it is a mechanical defense against chewing. However, they do not control for other factors that could also explain the observed data (such as higher plant investment in secondary compounds during folded/rolled stage). Grubb and coworkers (2008) hypothesize that leaf folding makes leaves harder to eat for many herbivores, and that this is one of the factors causing the higher rate of leaf damage by herbivores in dicots compared to monocots in tropical lowland rain forests that they observed. This is because folding/rolling of young leaves is a common monocot strategy and a far less common dicot strategy (Grubb et al., 2008).

There is not sufficient evidence in existing literature to claim that the nyctinastic folding of *A. pintoi* leaves is a defense against herbivory. In this study, the open and closed configurations of *A. pintoi* were tested through the construction of artificial leaves to see if one configuration reduces herbivory by *C. trogon*.

## METHODS

This experiment was performed in Monteverde, Costa Rica. *C. trogon* grasshoppers (Fig. 2a) were selected as the target herbivore for the experiment because of the ease of capturing a large number of them in the Monteverde area. The *Chromacris* grasshoppers naturally grouped together, likely for aposematic purposes (Despland, 2020), so 65 of them could be collected at once. *Chromacris* grasshoppers also performed well in laboratory conditions – they were easy to work with and ate a lot in petri dishes. *A. pintoi* leaves were not used; instead, artificial *A. pintoi* shaped leaves were constructed using *Iochroma arborescens* (known as güitite) leaves. *C. trogon* is not a natural herbivore of *A. pintoi*, so this study is only experimentally testing the plausibility of *A. pintoi* leaf folding being a herbivore defense, not confirming the adaptive significance of this behavior in a specific ecological setting.

**Figure 2.**
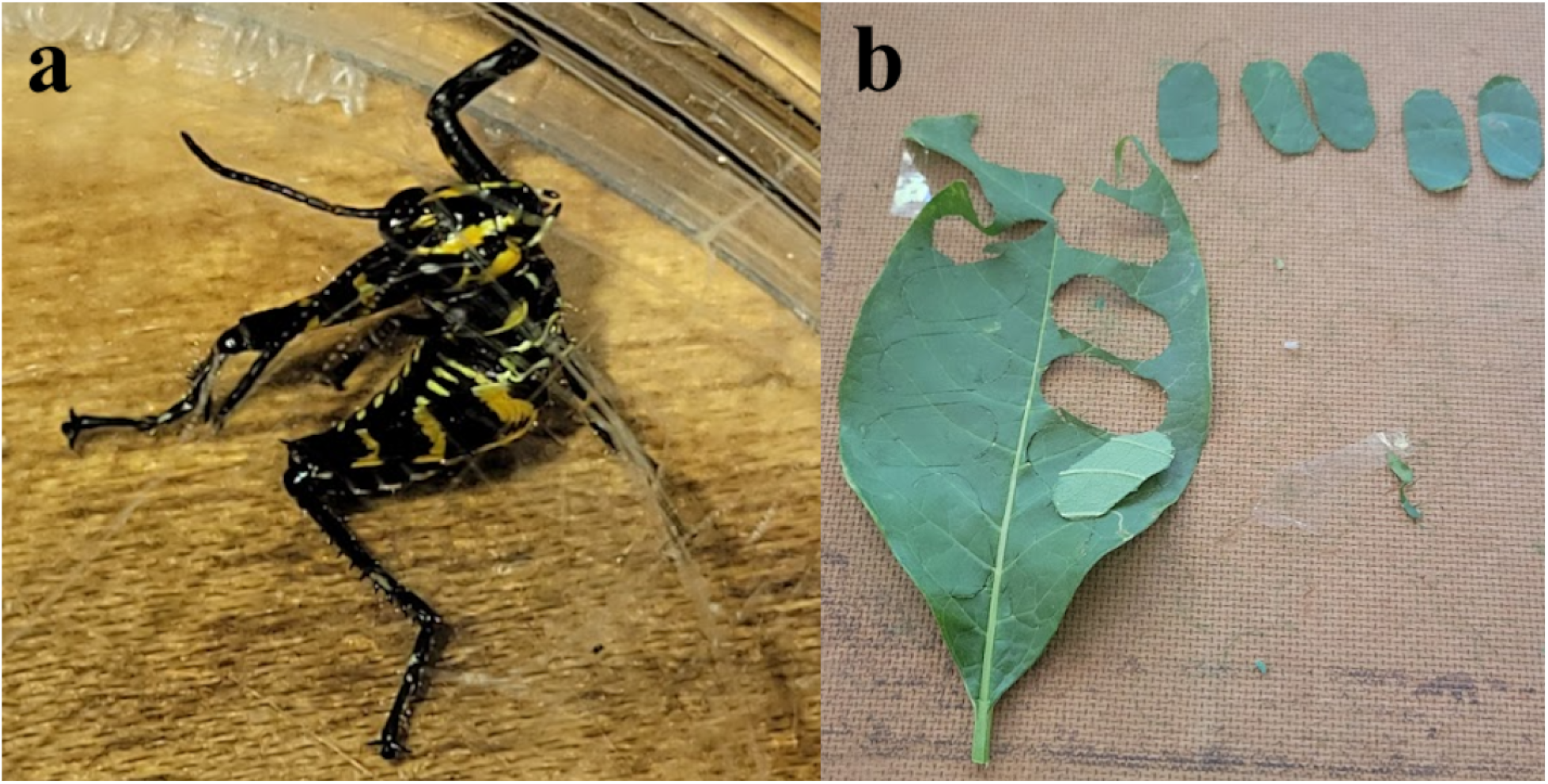
**a** *C. trogon* grasshopper used in the experiment. **b** Young *I. arborescens* leaf with leaflet cut-outs in it.

I constructed a leaflet cut-out tool using a thick stick, two scoopulas, duct tape, two razor blades, and two foldback metal clips. This tool acted like an oval cookie cutter, enabling me to cut out leaflets of the same size. Young *I. arborescens* stems with many leaves were broken off of trees the same day of an experimental trial, and their stems were placed in water prior to being processed into artificial leaves. 0.7 cm × 2 cm rounded leaflets were cut out of *I. arborescens* leaves using the leaf cutting tool (Fig. 2b). Based on a random sample of 30 leaflet pairs from the raw experimental data (15 closed, 15 open), the average percent difference between the area of any two leaflets was 7.8%±4.6%.

To construct open leaves, a 4 cm piece of dry grass was placed in the center of a 0.8cm × 2cm piece of duct tape, and leaflets were placed on tape following the template in Fig 3a. Closed leaves were constructed by placing dry grass of the same size on one side of an identically sized piece of tape (Fig. 3b), and placing leaflets onto the tape as in Figure 3a, 3b, and finally 3c. Closed leaves were constructed with the stomata side exposed on the top and bottom of the folded leaflet, like how *A. pintoi* folds (Fig. 1a). Even though a larger area of the closed leaf configuration is above the tape, tape is adhered to the same total amount of leaf surface area in both leaf types, because closed leaves overlap one another above the tape (Fig. 3). If leaflet stuck to tape is less desirable, this is controlled for in the experiment. However, if leaflet area *near* tape is less desirable for *C. trogon*, there may be a slight bias against closed leaves. Based on construction templates, 11.7% of open leaf area is directly above tape, while 19.5% of closed leaf area is directly above tape. Grasshoppers were observed to have eaten parts of leaflets directly stuck to tape from underneath twice, suggesting that tape was not a complete deterrent. I doubt that this small difference in leaf area near tape could have caused the strong effect observed, but this is a factor other than leaf shape that could have impacted results that must be acknowledged.

**Figure 3.**
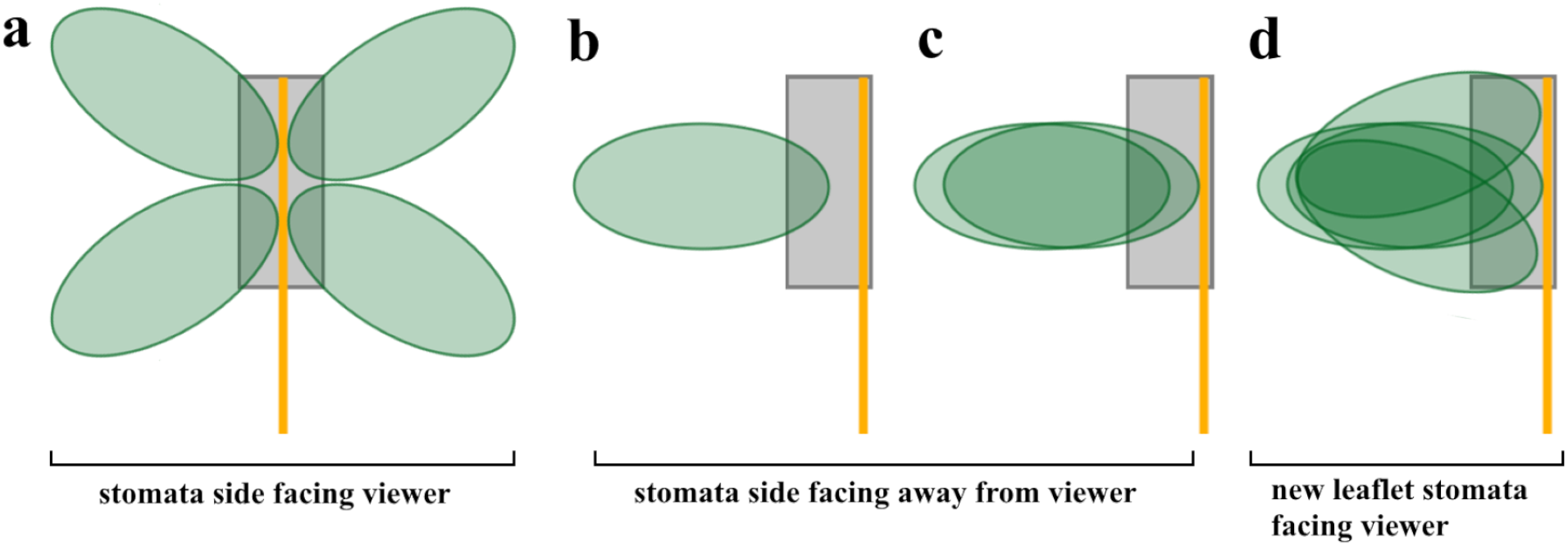
**a** Open artificial leaf template. **b**,**c**,**d** Closed leaf construction template. **b** First leaflet placed in closed leaf construction, stomata side down. **c** New leaf is the second leaflet placed in closed configuration, stomata side down. **d** Two new leaflets are the last two leaflets placed in closed leaf construction, with stomata sides up.

To mimic natural horizontal leaf posture, one closed and one open artificial leaf were taped to the top of petri dishes (Fig. 4a). While most species exhibiting FN assume a vertical position at night, *A. pintoi* often folds leaves into the ground, assuming a horizontal position (see Fig. 1a). Artificial petioles were bent such that part of all leaflets always touched the bottom of the petri dish. This allowed insects to go underneath the leaves if desired, mirroring natural conditions. Grasshoppers were selected from a pool of 65 captured *C. trogon*, and placed in petri dishes with open and closed leaves in groups of 3 (Fig. 4a). The experiment was performed during 3 separate days (6 trials, 17 trials, 7 trials), with random reshuffling of grasshoppers into new groups of 3 in between. All trials happened after a 24-hour period of starvation. Herbivory was measured after a six hour eating period.

**Figure 4.**
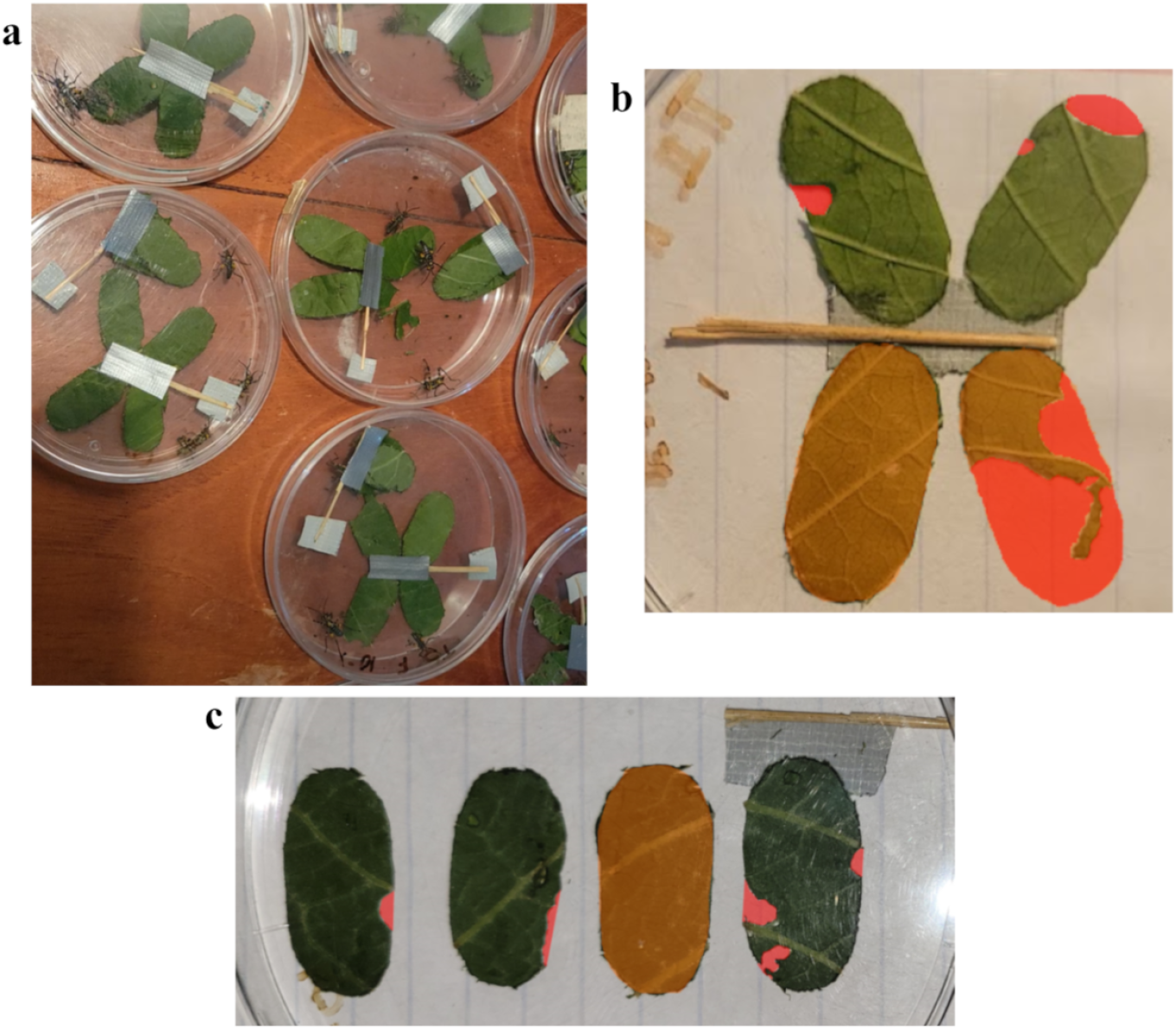
**a** Open and closed artificial leaves in petri dishes being eaten by *C. trogon* grasshoppers. **b** Picture of open and **c** closed artificial leaves after herbivory experiment, with eaten areas highlighted in red. See Appendix 2 for annotated images of herbivory areas in all 30 trials.

Herbivory percentage was calculated visually from pictures using area selections in the software paint.net. Usually, it was easy to manually select the eaten area (Fig. 4c), but sometimes there had been so much herbivory that the bounding edge of the leaflet had disappeared. In these cases, a reference leaflet selection was copied over and rotated to act as an overlay to guide selecting the other eaten leaflet area (Fig. 4b). Individual consumed pixel areas were summed and then divided by 4 times the area of the outline of the most complete leaflet (the orange leaves with no red in Fig. 4b,c) to determine the percentage of leaf area eaten for the whole leaf. A one tailed, paired t-test was performed on the open and closed leaf percent herbivory. The null hypothesis was that mean open leaf herbivory is not greater than mean closed leaf herbivory, and the alternative hypothesis was that mean open leaf herbivory is greater than mean closed leaf herbivory.

## RESULTS

Average herbivory of open leaves was 12.3%±6.9%, while the average herbivory of closed leaves was 5.2%±6.2% (Fig. 5). The difference between the average open herbivory and the average closed herbivory was 7.1%±11.9% (For raw data, see Appendix 1). A one-tailed t-test on the differences between open and closed herbivory yielded a p-value of 0.0014, indicating a significant preference for open leaves. Because 13 of the trials were randomized resamplings of the grasshoppers of the main 17 trials, I am assuming that the variability in day-to-day eating habits of grasshoppers and new groupings of three grasshoppers are enough to make the 13 trials independent of the main 17 trials. When the main 17 trials are analyzed alone, the p-value is 0.052.

**Figure 5.**
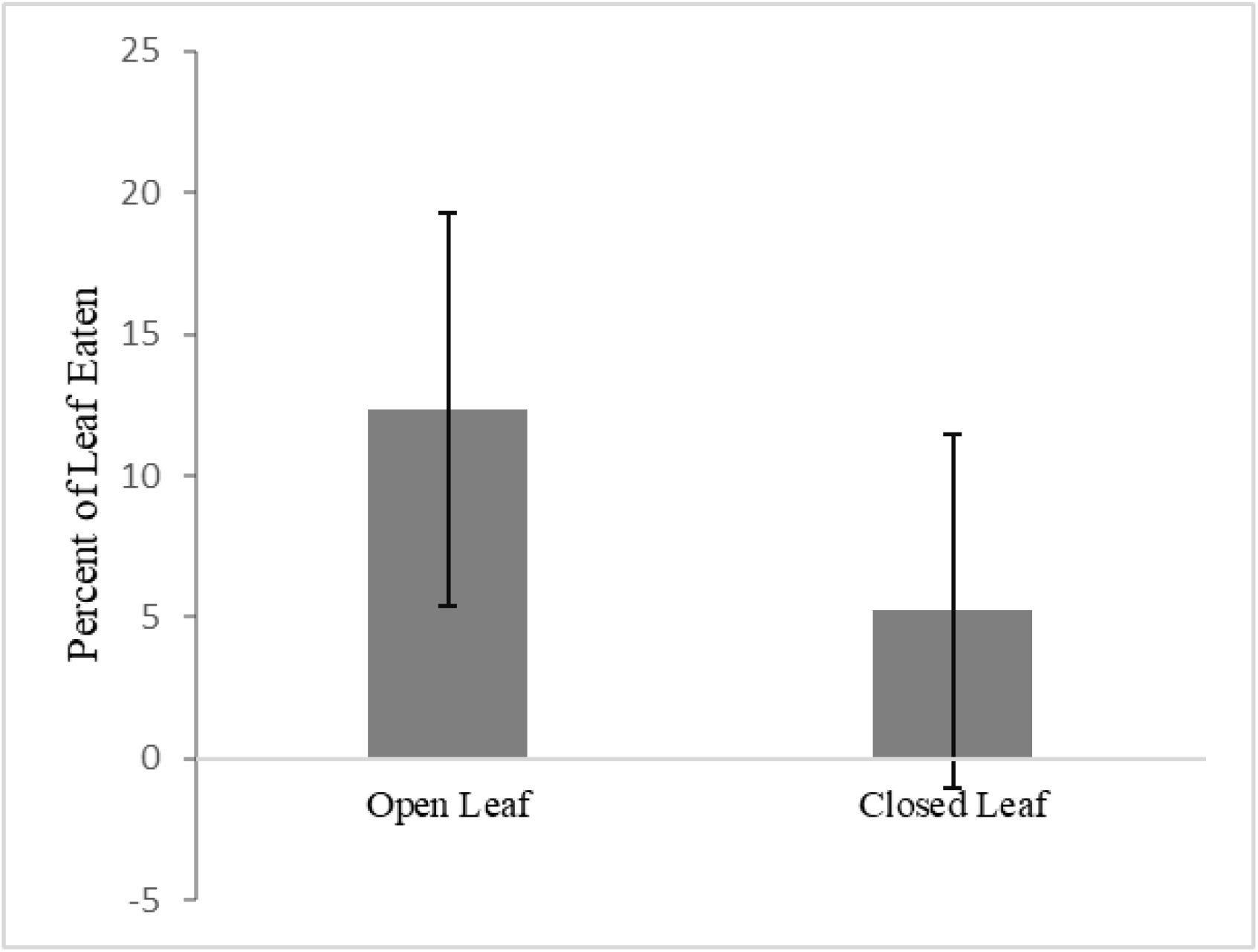
Average proportion of open and closed leaves eaten by 3 grasshoppers after 6 hours, error lines are standard deviation

Open leaves were eaten 2.3 times more than closed leaves by *C. trogon* grasshoppers. This result is strong evidence that with at least one herbivore, *A. pintoi* FN leaf folding acts as a defense against herbivory – supporting the hypothesis that an adaptive significance of leaf folding in *A. pintoi* is defense against herbivory.

## DISCUSSION

There are two main hypotheses that could explain the results of this experiment: (1) leaf folding is ‘camouflage’, reducing plant apparency (Bose, 1926; Minorsky, 2019) (2) leaf folding essentially makes leaves thicker and harder to eat (Grubb et al., 2008; Jackson et al., 1999).

Castagneyrol and coworkers (2013) demonstrated that plant apparency (defined as the likelihood of being found by herbivores, relative to alternative options) is significantly positively correlated with herbivory. The folded artificial leaf was certainly less apparent than the open leaf, as it had less exposed surface area, and took up less space in the petri dish. In the field, *A. pintoi* folds its four leaflets to look like one and further folds the petiole closer to the ground (Fig. 1), below other grass species if they are present, reducing apparency. The observed results could be because of less exposed surface area on closed leaves – if grasshoppers randomly ate exposed leaf surface area, they would end up eating more of the open leaves. Assuming random eating of exposed surface area by grasshoppers, the grasshoppers would have to actively prefer closed leaf surface area to have equal rates of herbivory between open and closed leaves.

Leaf folding making leaves harder to eat is the core of Grubb and coworkers’ (2008) hypothesis of the adaptive significance of leaf folding in reducing herbivory. Jackson and coworkers (1999) found that *Ectropis bistortat* caterpillars have lower rates of survival and growth when only fed young folded *Rosa canina* leaves compared with older unfolded leaves. If we assume that this result was because of the leaf folding, and not because of another factor such as a difference in investment in secondary compounds in new/older leaves, leaf folding must be making the leaf harder to eat, because apparency is irrelevant when there is only one option in a confined area. Additionally, across five species, Jackson and coworkers observed that the most damage in new folded or rolled leaves in the field was caused by galling and boring insects that are not affected by stacked leaves increasing thickness. Folding mechanically making leaf tissue harder to eat can explain the results of this artificial leaf study. However, it must be noted that the cutout leaflets were very floppy and were not strongly pressed against one another, so the folding in this study did not increase leaf toughness as much as it does in other plants, e.g. in young Marantaceae leaves. It is unclear whether decreased apparency or increased toughness were the primary explanation for the observed data. Further research needs to be done to isolate the effects of apparency and folding making leaves harder to eat.

In a study by Perez-Harguindeguy and coworkers (2008), it was demonstrated that when leaves are not specifically accessible or specifically consumed, there is a high correlation between generalist preference in herbivory experiments and leaf consumption in the field. Because this experiment was done using the same leaf substrate for both conditions, the substrate was acceptable to the target species, and there was no difference in accessibility (other than being folded vs. open), it is reasonable to assume that the results of this herbivore preference experiment are correlated with herbivory in the field.

This study is evidence that folding can be a herbivore defense, but it is important to note that herbivores have co-evolved with folded plant leaves. Hispine beetle larvae are flattened to fit in between tight layers of palm or ginger leaves (Strong, 1977; McKenna & Farrell, 2005). Similar to how sometimes secondary compounds actually become beneficial to herbivores (Smilanich et al., 2016), it is likely that in other plant-herbivore interactions, leaf folding is actually attractive to herbivores. More leaf folding herbivore preference experiments need to be done with different herbivores and different folding leaf types, and using the actual species that are nyctinastic.

## ACKNOWLEDGEMENTS

I am so grateful to have had the mentorship of Emilia Triana and Naomi Solano in this project. Their wisdom, combined with the accessibility of EAP resources and materials, made this project possible. Thank you to Mauricio Vargas of Finca la Florencia for showing me *C. trogon* on a leaf in his farm.

