## Appendix for "Nyctinastic leaf folding mimic reduces herbivory by *Chromacris trogon* grasshoppers (Orthoptera:Romaleidae)"

**Appendix 1** | Leaf proportion eaten per trial:

| | Open Leaf Proportion Eaten | Closed Leaf Proportion Eaten | Difference of open and closed |
|---|---|---|---|
| Trial 1 | 0.17019329 | 0 | 0.170193 |
| Trial 2 | 0.17701515 | 0.03340218 | 0.143613 |
| Trial 3 | 0.01750776 | 0.16625524 | -0.14875 |
| Trial 4 | 0.17155656 | 0 | 0.171557 |
| Trial 5 | 0.18886345 | 0.01225531 | 0.176608 |
| Trial 6 | 0.19019648 | 0 | 0.190196 |
| Trial 7 | 0.02208923 | 0.07828447 | -0.0562 |
| Trial 8 | 0.13165118 | 0.02519099 | 0.10646 |
| Trial 9 | 0.1326294 | 0.06891748 | 0.063712 |
| Trial 10 | 0.07437332 | 0.15502927 | -0.08066 |
| Trial 11 | 0.18139945 | 0.06877005 | 0.112629 |
| Trial 12 | 0.10226598 | 0 | 0.102266 |
| Trial 13 | 0.05093532 | 0.07083841 | -0.0199 |
| Trial 14 | 0.20439722 | 0 | 0.204397 |
| Trial 15 | 0 | 0.25169432 | -0.25169 |
| Trial 16 | 0.04089851 | 0.01513947 | 0.025759 |
| Trial 17 | 0.05415896 | 0.03897076 | 0.015188 |
| Trial 18 | 0.25819524 | 0 | 0.258195 |
| Trial 19 | 0.22130354 | 0.02706651 | 0.194237 |
| Trial 20 | 0.20917896 | 0.00882875 | 0.20035 |
| Trial 21 | 0.07577493 | 0 | 0.075775 |
| Trial 22 | 0.09724929 | 0.10078256 | -0.00353 |
| Trial 23 | 0.083325 | 0.09164315 | -0.00832 |
| Trial 24 | 0.10950638 | 0.14125884 | -0.03175 |
| Trial 25 | 0.14456528 | 0.05320266 | 0.091363 |
| Trial 26 | 0.20093145 | 0 | 0.200931 |
| Trial 27 | 0.1384412 | 0 | 0.138441 |
| Trial 28 | 0.12269312 | 0.04571073 | 0.076982 |
| Trial 29 | 0.11197726 | 0 | 0.111977 |
| Trial 30 | 0.0194377 | 0.11136889 | -0.09193 |

**Appendix 2** | Raw unanalyzed photos of open and closed leaves after the experiment:
https://photos.app.goo.gl/u4iNVnq3bPeR4XXi6

All analyzed leaves with herbivory highlighted in red (leaves with no area eaten not included):
https://photos.app.goo.gl/79Vui7aYn8GMbYwW7
